# Rapid structural remodeling of peripheral taste fibers is independent of taste cell turnover

**DOI:** 10.1101/2022.11.17.513266

**Authors:** Zachary D. Whiddon, Jaleia Marshall, David Alston, Aaron W. McGee, Robin F. Krimm

**Affiliations:** Department of Anatomical Sciences and Neurobiology, University of Louisville School of Medicine, Louisville, KY 40202

**Keywords:** taste, geniculate neuron, morphology, afferent, tongue

## Abstract

Taste bud cells are constantly replaced in taste buds as old cells die and new cells migrate into the bud. The perception of taste relies on new taste bud cells integrating with existing neural circuitry, yet how these new cells connect with a taste ganglion neuron is unknown. Do taste ganglion neurons remodel to accommodate taste bud cell renewal? If so, how much of the taste axon structure is fixed and how much remodels? Here we measured the motility and branching of individual taste arbors (the portion of the axon innervating taste buds) over time with two-photon *in vivo* microscopy. Terminal branches of taste arbors continuously and rapidly remodel within the taste bud. This remodeling is faster than predicted by taste bud cell renewal, with terminal branches added and lost concurrently. Surprisingly, ablating new taste cells with chemotherapeutic agents revealed that remodeling of the terminal branches of taste arbors does not rely of the renewal of taste bud cells. Although the arbor structure remodeling was fast and intrinsically controlled, no new arbors were added, and few were lost over 100 days. Taste ganglion neurons maintain a stable number of nerve arbors that are each capable of high-speed remodeling. Arbor structural plasticity would permit arbors to locate new taste bud cells, while stability of arbor number could support constancy in the degree of connectivity and function for each neuron over time.

## Introduction

The replacement of cells in the taste bud was discovered more than fifty years ago (Beidler and Smallman, 1965). In fact, cells within a taste bud are one of the fastest renewing cell populations in the body, with an average lifespan of 10 days (Beidler and Smallman, 1965; Farbman, 1980; Lindemann, 2001). Taste bud cells transduce chemical stimuli in the oral cavity, and transmit a signal to the axons of taste ganglion neurons, which carry this information to the brain. Each time a taste bud cell dies, the axon of a taste ganglion neuron must connect with a new taste bud cell if it is to maintain functional integrity (Hirose et al., 2020; Kumari et al., 2017; Lee et al., 2017). How this process is orchestrated is still unclear, but could involve structural remodeling of the neuron (Donnelly et al., 2022; Zaidi et al., 2016). Alternatively, taste bud cells could migrate to the axon of the taste ganglion neuron, making neuronal remodeling unnecessary.

Progress toward understanding the cellular movements occurring during taste bud cell turnover has been hindered by the use of fixed-tissue analysis (Beidler and Smallman, 1965; Farbman, 1980; Kinnamon et al., 1988; Wilson et al., 2022). Because cells are chemically fixed into position for observation, this approach fails to capture the dynamic processes within the taste bud, such as taste cell movements and axon remodeling. (Beidler and Smallman, 1965; Farbman, 1980; Ohman et al., in submission). Fixed-tissue studies also fail to describe what happens to axon remodeling when cell turnover is disrupted (e.g. chemotherapies) (Donnelly et al., 2022; Kumari et al., 2017, 2015). Additionally, because it is unknown whether taste ganglion neurons undergo structural remodeling, it is unclear why there are morphological differences between individual taste ganglion neurons (Huang et al., 2021). Morphological differences between individual taste ganglion neurons could be inherent features of the neuron that dictate differences in functional properties or could be snap-shots in time of a continuously remodeling circuit that lacks functional stability. Understanding how taste ganglion neuron structure changes over time is a fundamental step in relating taste ganglion neuron morphology to function in the taste system.

In other parts of the nervous system, longitudinal *in vivo* imaging has been used to study the structural plasticity of axons and dendrites over time. Structural remodeling has been shown to occur during development and following injury or disease (Gangadharan et al., 2022; Martineau et al., 2018; Nakazawa et al., 2018; Speidel, 1935, 1933, 1932; Walsh and Lichtman, 2003). Under these conditions, axonal degeneration is the process that has been most extensively described (Bernstein and Lichtman, 1999; Bishop et al., 2004). In adult neurons, dendritic branches are mostly stable and remodeling is restricted to dendritic spine turnover (Grutzendler et al., 2002; Purves and Hadley, 1985; Trachtenberg et al., 2002). Axons may show a greater tendency to remodel than dendrites, but this remodeling is neuron type specific (De Paola et al., 2006). In the taste system, it has not been determined whether peripheral axons have the capacity remodel and to what extent remodeling occurs.

Here we examined the structure of peripheral taste axons using *in vivo* two-photon microscopy to determine how taste ganglion neuron structure changes over time. We discovered that peripheral axons of taste ganglion neurons have regions that rapidly remodel and regions that are stable throughout adulthood. Terminal branches remodel rapidly and axons undergo sufficient levels of remodeling to connect with new taste bud cells over time. The structural plasticity within taste ganglion neurons is an inherent feature of the neurons and is not regulated by factors from the taste bud cells. Structural stability maintains total number of axonal arbors for each taste ganglion neuron, and this stability supports consistency in the total number of taste-transducing cells providing input to an individual taste ganglion neuron. We conclude that taste ganglion neurons maintain their peripheral circuits by having a stable number of axonal arbors, with each arbor continuously remodeling to connect with new taste bud cells over time.

## Results

### Acute *in vivo* imaging reveals axonal structural plasticity

To determine whether axon structure in the peripheral taste system is fixed or remodels over time, we developed an *in vivo* preparation to image the portion of the axon of a taste ganglion neuron that terminates in the taste bud (axonal arbors). The arbors of taste ganglion neurons enter taste buds in the epithelium. On the front two-thirds of the mammalian tongue, taste buds are housed in epithelial structures called fungiform papillae, which are easily accessible for imaging in live animals (Miller and Reedy, 1990). Within taste buds, arbors form a dense plexus with many overlapping branches (Ohman-Gault et al., 2017; Wilson et al., 2022). We used a sparse-labeling strategy to express tdTomato in a small number of taste ganglion neurons so that individual arbors could be distinguished (Huang et al., 2021). In this preparation, fewer than five labeled arbors were labeled with tdTomato per fungiform taste bud and many taste buds did not contain any labeled arbors. Individual arbors were imaged using two-photon laser scanning microscopy (Figure 1A).

**Figure 1.**
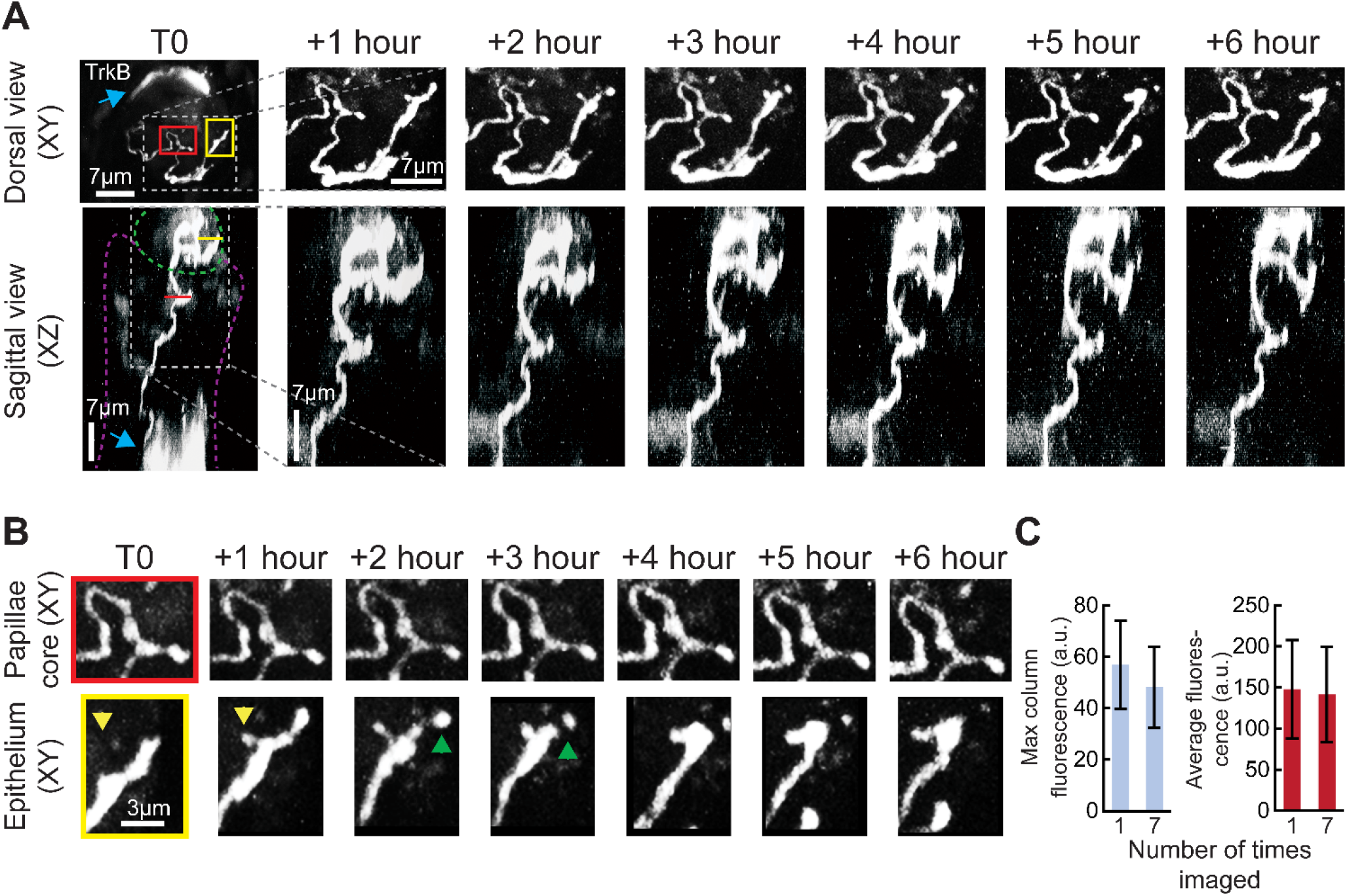
Intravital imaging of taste neurons in fungiform papillae. Example of a fungiform papilla with a single labeled receptive arbor illustrating the small-scale changes that can be seen within 6 hours. (**A**) Top and bottom rows show dorsal (XY) and sagittal (XZ) views of an arbor within a fungiform papilla every hour over a 6 h period. Some keratinocytes were also labeled (Liu et al., 2012; Tang et al., 2017) (cyan arrow). After T0, images for each consecutive timepoint are enlarged to show only the labeled arbor. At T0, the taste bud is outlined in a green dotted line, and the connective tissue core of the papilla is outlined in magenta. (**B**) High-magnification view (XY projection) of structural features of the arbor within the papilla core (red box) and taste bud (yellow box). The branch ending inside the taste bud bifurcated after 1 h and then reverted to a single branch ending after 4 h (yellow and green arrowheads). The portion of the arbor in the papilla core did not change across 6 h. (**C**) Average and maximum fluorescence (mean ±SD), using regions of interest around receptive arbors, in max intensity z-projections of arbors imaged every hour for 6 h. Fluorescence levels between hour 1 and 6 are unchanged.

Individual arbors were imaged up to a depth of 70 microns to capture the entire arbor, as well as a portion of the axon beneath the taste bud. To estimate the frequency of structural plasticity for these axons, first we imaged the same arbors once per hour for a 6 h period (n=36). We found that arbor structure within the lamina propria (Figure 1A, purple dotted-line) of the fungiform papilla core was stable (Figure 1B, yellow box). Many of the terminal branches of arbors within the taste bud (Figure 1A, green dotted line) displayed structural changes in as quickly as a few hours after the initial imaging session (Figure 1B, red box). Among the 36 arbors imaged, 24 changed their number of terminal branches in a 6 h window by adding and/or subtracting small terminal branches (Figure 1B, yellow box).

To determine whether the very low excitation power (<51 mW) used for imaging induced evident photobleaching, we imaged a new group of arbors (n=13 from two mice) using identical acquisition parameters every hour for 6 h and compared the fluorescence intensity across image stacks (z-max intensity projections) for the first and last timepoints. We did not observe a significant decrease in average (t(12) = −1.78, p = 0.076) or maximum (t(12) = −1.78, p = 0.380) fluorescence intensity despite imaging these structures repeatedly (Figure 1C). Thus, this imaging approach was suitable for acute *in vivo* imaging of individual axonal arbors and demonstrates that arbors of taste ganglion neurons display high-speed structural remodeling.

### Chronic *in vivo* imaging requires a fungiform papillae map

Next, we examined axonal structural plasticity over days. Reliably identifying the same taste buds across imaging sessions was critical for chronic imaging experiments. To identify the same taste buds over days or weeks, we combined our approach for sparse labeling of taste ganglion neurons (*Ntrk2*^CreER^ :tdTomato) with transgenic mice that express green fluorescent protein (GFP) in taste buds (*Sox2*^GFP^: Okubo et al., 2006). When viewed at low-magnification, GFP expression on the anterior tongue surface revealed a unique taste bud pattern for each mouse (Figure 2A) This provided a set of fiduciary marks used to locate the same taste buds and arbors repeatedly over time. Three taste buds in the example papillae map contained labeled arbors; they are shown at high magnification 24 h apart (Figure 2B). No other fungiform taste buds viewed in this field contained labeled arbors because of the sparseness of genetic labeling of taste ganglion neurons (Figure 2A, taste buds with no box).

**Figure 2.**
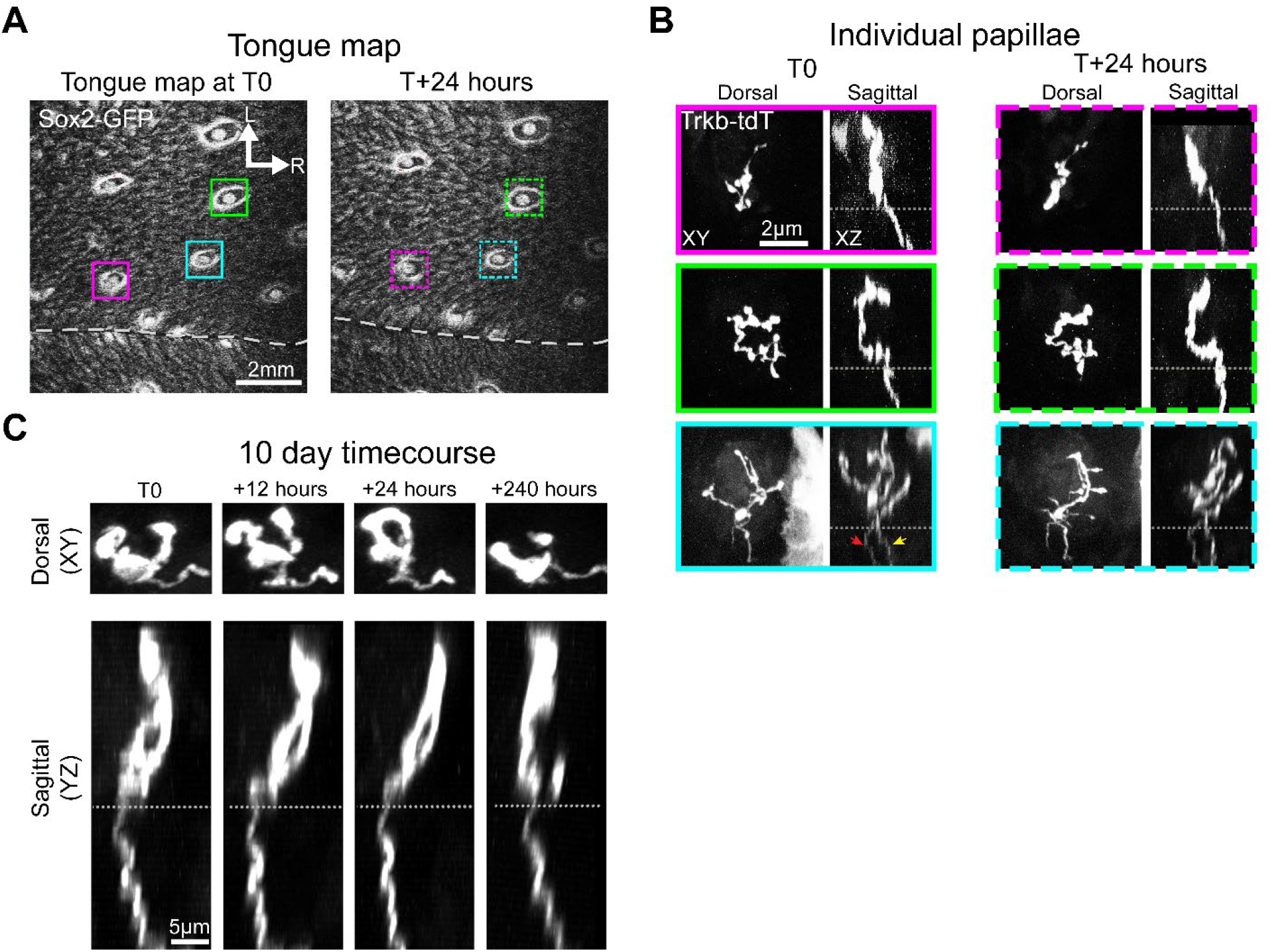
Chronic intravital imaging using a papillae map. (**A**) Sox2-GFP expression in the anterior tongue epithelium illustrating a papillae map. Papillae with labeled arbors are indicated by solid boxes at T0 and dashed boxes 24 h later. The tongue midline is indicated with a white dashed line, and rostral (R) and lateral (L) directions are indicated by white arrows. (**B**) High-magnification micrographs of TrkB-tdTomato-labeled nerve arbors in each of the three papillae indicated from the map. The first column shows the XY (dorsal) plane, which is parallel to the tongue surface, and the XZ plane is shown in the adjacent column. Gray dashed lines in the z-plane images illustrate the bottom of the taste bud. The papillae outlined in magenta and green have a single labeled arbor, and the papilla outlined in cyan has two separate labeled arbors (yellow and red arrows). Micrographs outlined in dashed lines show high-magnification views of the same papillae 24 h later. Some changes to each arbor structure can be seen during this time within the taste bud but not within the papillae core. (**C**) Receptive arbors were imaged every 12 h for 10 days, and an example arbor is shown at the initial timepoint (T0) as well as 12, 24, and 240 h later.

Taste bud cells have an average lifespan of 10 days (Beidler and Smallman, 1965). Given this rate of taste bud cell turnover, we investigated the structural plasticity of arbors every 12 h for 10 days. In five mice, we observed the same 31 taste buds and 60 labeled arbors 21 times over 10 days. Among these arbors, four were lost during this period, and none were added. One taste bud lost GFP expression in the epithelium, but the arbor remained, consistent with taste bud loss due to aging (Liebl et al., 1999; Patel and Krimm, 2010). Examination of a single-labeled arbor at the beginning and end of the 10-day experiment illustrates that arbor structure in the papillae core was stable, whereas arbor structure in the epithelium was plastic (Figure 2C).

### Terminal branches remodel rapidly and continuously over 10 days

The axonal arbors of taste ganglion neurons synapse with taste bud cells (Kinnamon et al., 1988; Romanov et al., 2018; Vandenbeuch et al., 2015; Wilson et al., 2022). Many taste axonal arbors contact taste bud cells on terminal branches (Ohman et al., in submission). If the turnover of taste bud cells regulates arbor structure, then terminal branches are likely to remodel as taste bud cells die and/or new cells enter the taste bud. Arbors form synapses with an average of 1.6 taste bud cells (Kinnamon et al., 1988). If new terminal branches are added and lost as a means of synapsing with new taste bud cells, then most arbors should add or lose roughly one terminal branch over 10 days, and some arbors are likely to remain stable.

In five mice, we imaged 31 taste buds containing a total of 60 arbors. However, taste buds with three or more labeled arbors were not quantified because we could not distinguish arbors when branches intermingled. We identified 26 individual arbors from 21 different taste buds that were distinguishable at all timepoints and quantified their terminal branches every 12 h (Figure 3A-B, video 1). All quantified arbors added and lost multiple terminal branches during within 10 days (Figure 3B-C). In 43% of arbors, a terminal branch was lost or added within the first 12 h, and 100% of arbors gained or lost a branch within the first 4.5 days (108 h, Figure 3D). For the 30 arbors imaged every 12 h for at least 4 days, one terminal branch was added or lost every 16.8 ± 1.1 h. This anatomical plasticity was much faster than predicted based on the rate of taste bud cell turnover. Thus, terminal branches do not appear to be added for the primary purpose of synapsing with a new taste bud cell or lost solely as a mechanism for disconnecting from dying taste bud cells.

**Figure 3.**
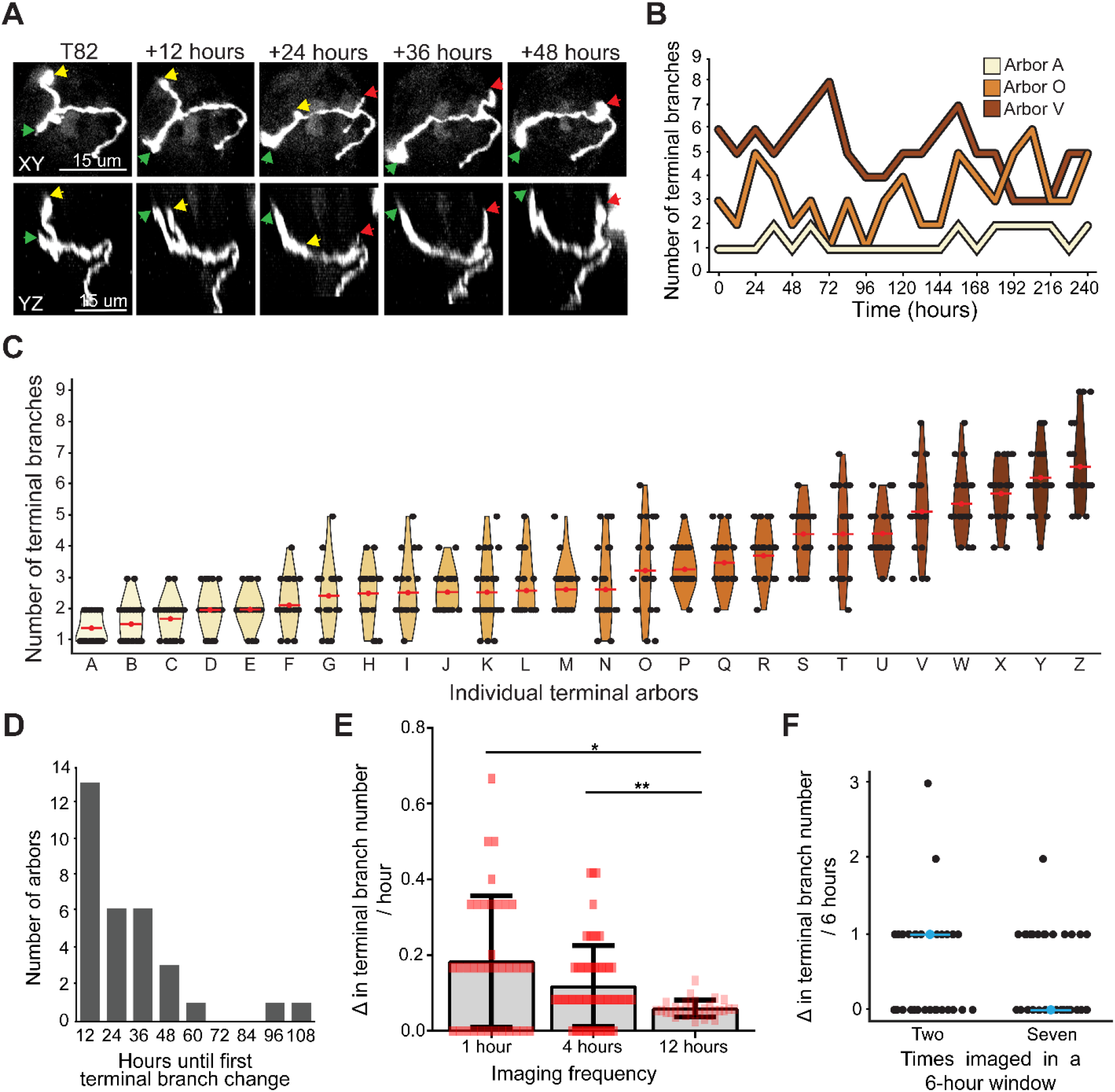
Terminal branches are continuously and rapidly remodeled. (**A**) Example arbor shown every 12 h over 2 days (for the full 10 days see Figure 3, see supplementary movie 1 for full 10 days). Terminal branches labeled with green and red arrowheads extended during this period, while the terminal branch labeled with a yellow arrowhead retracted. (**B**) Number of terminal branches every 12 h for three arbors across the 10-day imaging window. (**C**) Violin plots for 26 (A-Z) individual arbors showing the distribution of terminal branch number during the 10-day imaging window. Arbors are arranged in ascending order of average terminal branch number (red dot and bar) across 10 days. (**D**) Amount of time until the first gain or loss of a terminal branch. All arbors added or lost a terminal branch within 108 h. (**E**) Violin plots showing rates of change in terminal branch number per hour for three different sampling frequencies: 1, 4, or 12 h. Median values are indicated with cyan dots and bars (Single asterisk p<0.05, double p<0.001). (**F**) Rate of change in terminal branches over 6 h for arbors imaged twice compared with arbors imaged seven times. Median values are indicated with cyan dots and bars.

Given the high rate of terminal branch addition and removal, we suspected that we could be underestimating the rate of change if some terminal branches were both gained and lost within the 12 h intervals between imaging sessions. To test this possibility, we imaged every 4 h for a 12 h period. For this image set (n = 57 arbors), an average of one terminal branch was added or lost every 6.6 ± 1.0 h, which was a significantly faster rate of change than the one obtained when arbors were imaged every 12 h (16.8 h; Figure 3E, U = 470.5, p < 0.001, Mann-Whitney U). These results indicate that imaging every 12 h underestimates the rate of terminal branch change. To verify the accuracy of this rate of terminal branch change, we compared arbors imaged every 4 h to arbors imaged every hour for 6 h (n = 36) and found no significant difference in the rate of terminal branch gain/loss (1 h rate: 5.5 ± 0.8 h; 4 h rate, 6.6 ± 1.0 h; U = 822.5, p = 0.09) (Figure 3E). Thus, we conclude that accurately describing changes in terminal branch number requires sampling arbor structure every 4 h. However, given the high rate of terminal branch addition and loss, we also sought to determine whether two-photon exposure influences the rate of terminal branch addition/loss. We compared arbors imaged twice in 6 h with arbors imaged seven times over the same amount of time (Figure 3F). There was no difference in the rate of change in terminal branch number over the 6 h period between these two sets of arbors, indicating that image collection does not influence the rate at which terminal branches are added and lost.

### Gain/loss of terminal branches is not regulated by taste bud cell turnover

If arbors add new branches in order to synapse with new taste bud cells, then altering the rate of new cell entry would be expected to alter terminal branch dynamics. To test the hypothesis that taste ganglion neuron remodeling is dependent on the addition of new taste bud cells we prevented new cells from being added to taste buds by blocking Hh-signaling. Administering LDE225 (trade name Sonidegib) (Figure 4A) stops new taste cells from differentiating and entering taste buds (Castillo et al., 2014; Castillo-Azofeifa et al., 2017; Ermilov et al., 2016; Kumari et al., 2017, 2015; Lu et al., 2018) (Figure S1). This is a chemotherapeutic agent used to treat basal cell carcinoma and is associated with loss of taste by patients receiving treatment (Jacobsen et al., 2016; Kumari et al., 2015; Lacouture et al., 2016; Tang et al., 2012).

**Figure 4.**
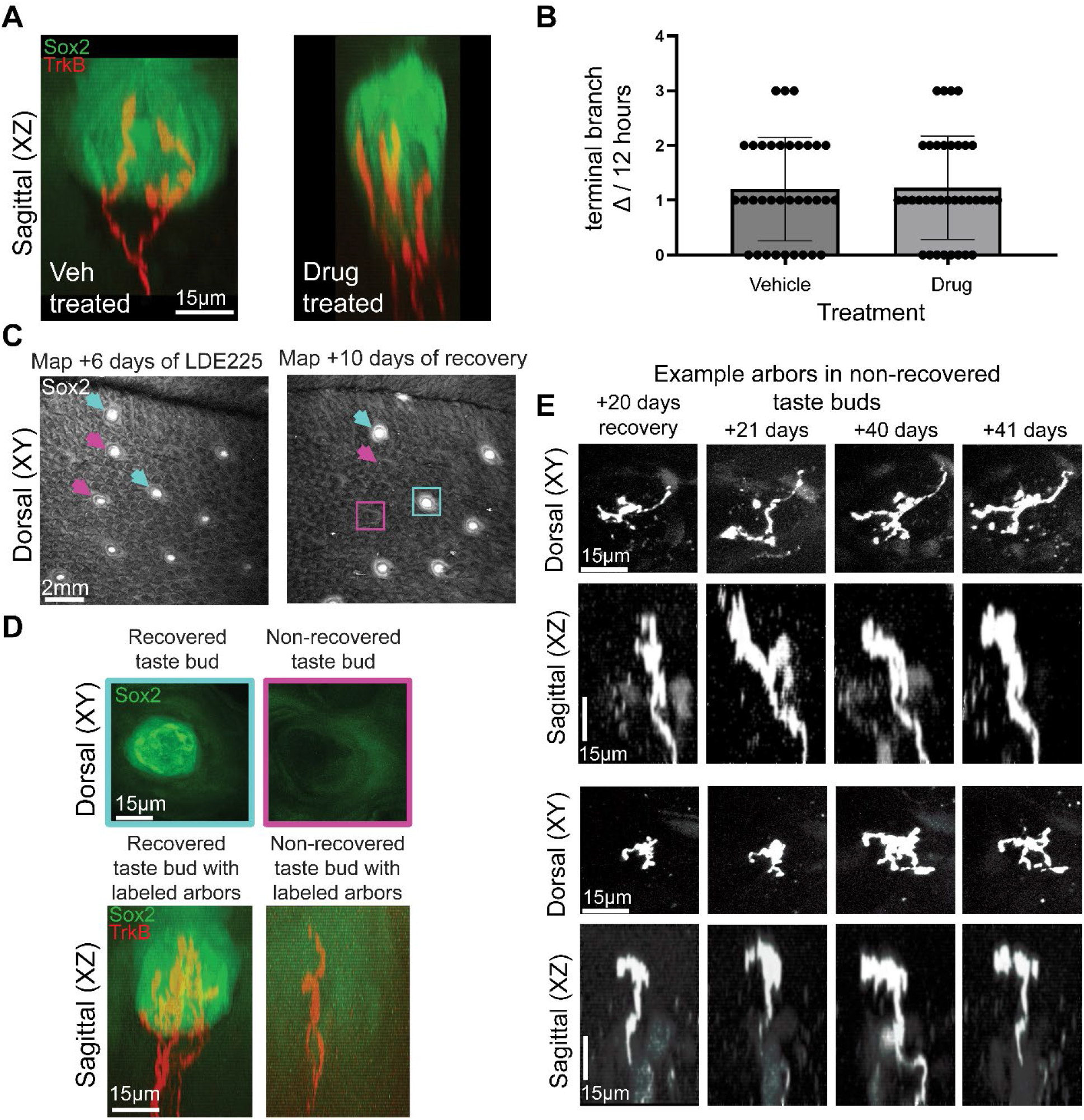
Terminal branch remodeling is not influenced by addition of new cells into the taste bud. (**A**) Example vehicle-treated and drug-treated taste buds after 8 days of treatment. (**B**) There was no difference in the rate with which terminal branches were gained/lost when taste bud cell turnover was halted (LDE225-treated mice) compared with normal conditions (vehicle-treated/untreated mice). (**C**) After LDE225 treatment, some fungiform papillae did not regain taste buds as determined by Sox2-GFP (magenta arrows and box). Two recovered taste buds are indicated with blue arrows and box. (**D**) Individual fungiform papillae with a taste bud had high levels of Sox2 expression, whereas papillae without a taste bud lacked cells with high levels of Sox2 expression. Papillae with both recovered and non-recovered taste buds retained taste ganglion neuron innervation. (**E**) Two papillae containing non-recovered taste buds with single-labeled arbors are shown 20, 21, 40, and 41 days after LDE225 treatment. Both arbors continued to remodel somewhat despite the absence of a taste bud.

**Figure 4 - Supplemental 1.**
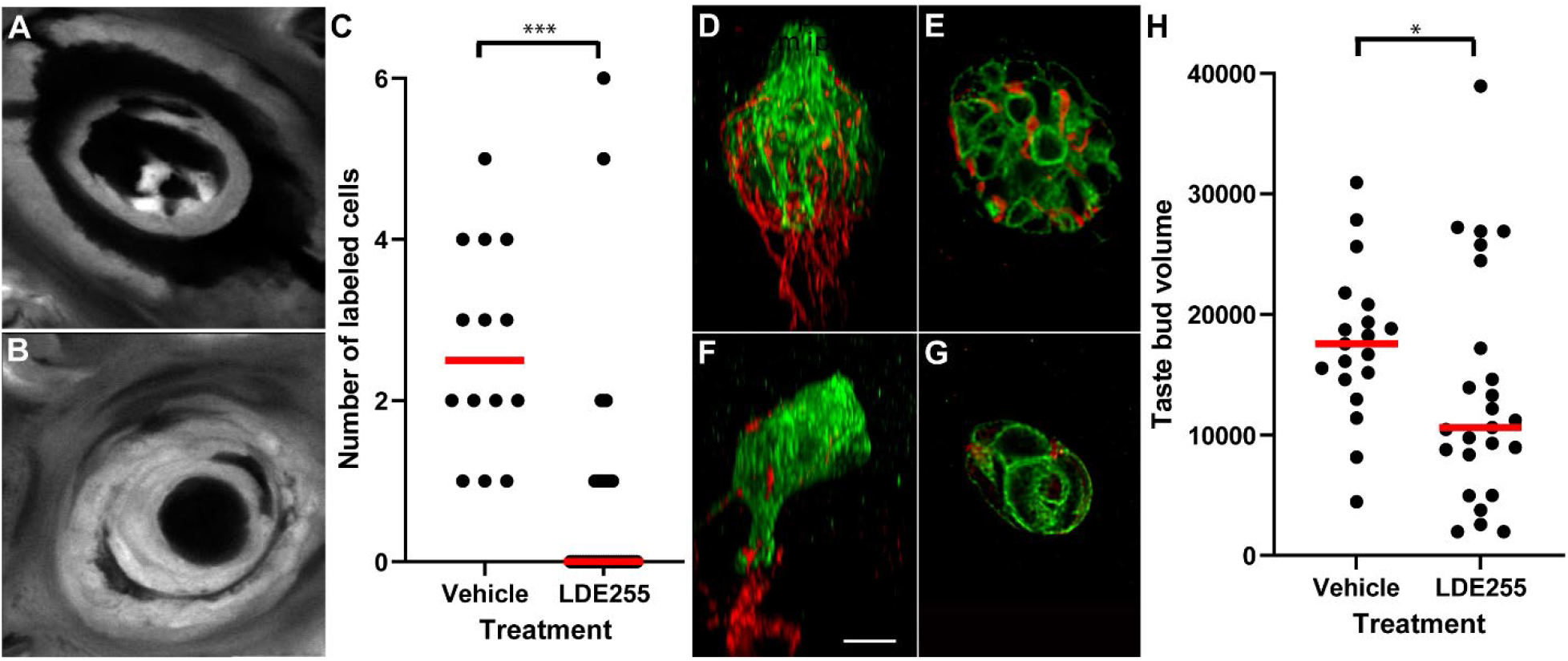
LDE225 inhibits new cell entry into the taste bud, resulting in volume reduction by day 10. **A-C)** A subset of epidermal precursor and/or stem cells expressing K14-CreER:tdTomato were labeled with a single dose of tamoxifen immediately preceding 10 days of treatment with vehicle (**A**) or LDE225 (**B**). On day 10, LDE255-treated mice had fewer labeled cells in taste buds than vehicle-treated mice (**C**). Specifically, 27 of 40 taste buds completely lacked new labeled cells following LDE255 treatment, while all those examined had some labeled cells following vehicle treatment. (**D-H**) Whole mounts of the lingual epithelium were labeled for taste buds (green) and neurons (Phox2b-tdTomato, red) following 10-day of treatment with vehicle (**D** z-plane, **E** x-y plane) and LDE225 (**F** z-plane, **G** x-y plane) tongues. (**H**) Taste bud volumes were smaller following 10 days of LDE225 compared with vehicle controls. *p ≤ 0.05, ***p ≤ 0.001.

We treated mice with LDE225 (n=4) or vehicle (n=2) for 10 days and imaged taste axonal arbors every 4 h for 12 h on days 6, 8, and 10. Over time, old taste cells continue to die and taste bud volume decreases (Castillo-Azofeifa et al., 2017) (Figure S1). However, arbors had similar rates of terminal branch change on each day of imaging (H(30) = 29.5, p = 0.44). Terminal branch gain/loss was similar between the LDE225 and vehicle groups (U = 312, p = 0.61) (Figure 4B). Thus, we conclude that preventing new cells from entering taste buds does not alter the rate of terminal branch remodeling.

We also examined how taste buds recover following treatment with LDE255. Many fungiform papillae recovered normal taste bud morphology, however some failed to recover and lacked Sox2 expression up to 40 days after treatment for 10 days with LDE225 (Figure 4C). Non-recovered papillae retained taste ganglion neuron innervation even in the absence of taste buds (Figure 4D). Surprisingly, we observed at least some remodeling of terminal branches in all papillae that lacked taste buds over 20-day, although remodeling over 24-hour was limited. Two example papillae with single labeled arbors are shown in Figure 4E. We conclude from these findings that the taste bud is not required for terminal branch remodeling of taste arbors, although remodeling may be slowed.

### Growth and retraction occur concurrently within the arbor

Some terminal branches displayed distinct morphologies as swellings on the tip of the terminal branch, as well as punctate swellings along the axon (Figure 5A). We suspected these features could be associated with remodeling, as both retraction bulbs and growth cones have been described in taste arbors (Zaidi et al., 2016). To determine if terminal branch swellings were associated with gain or loss of terminal branches, we examined 12 arbors from three mice that were imaged every 12 h for 10 days (n = 252 total timepoints). We observed 72 instances of swellings at the tip of the terminal branch and/or branch swellings long the branch. These terminal swellings along the axon were followed by retraction of the branch 86.1% of the time, whereas only 13.9% of terminal branches displayed terminal branch swelling without retracting (Figure 5B). Therefore, we conclude that terminal end swellings are often associated with retracting terminal branches (Figure 5A).

**Figure 5.**
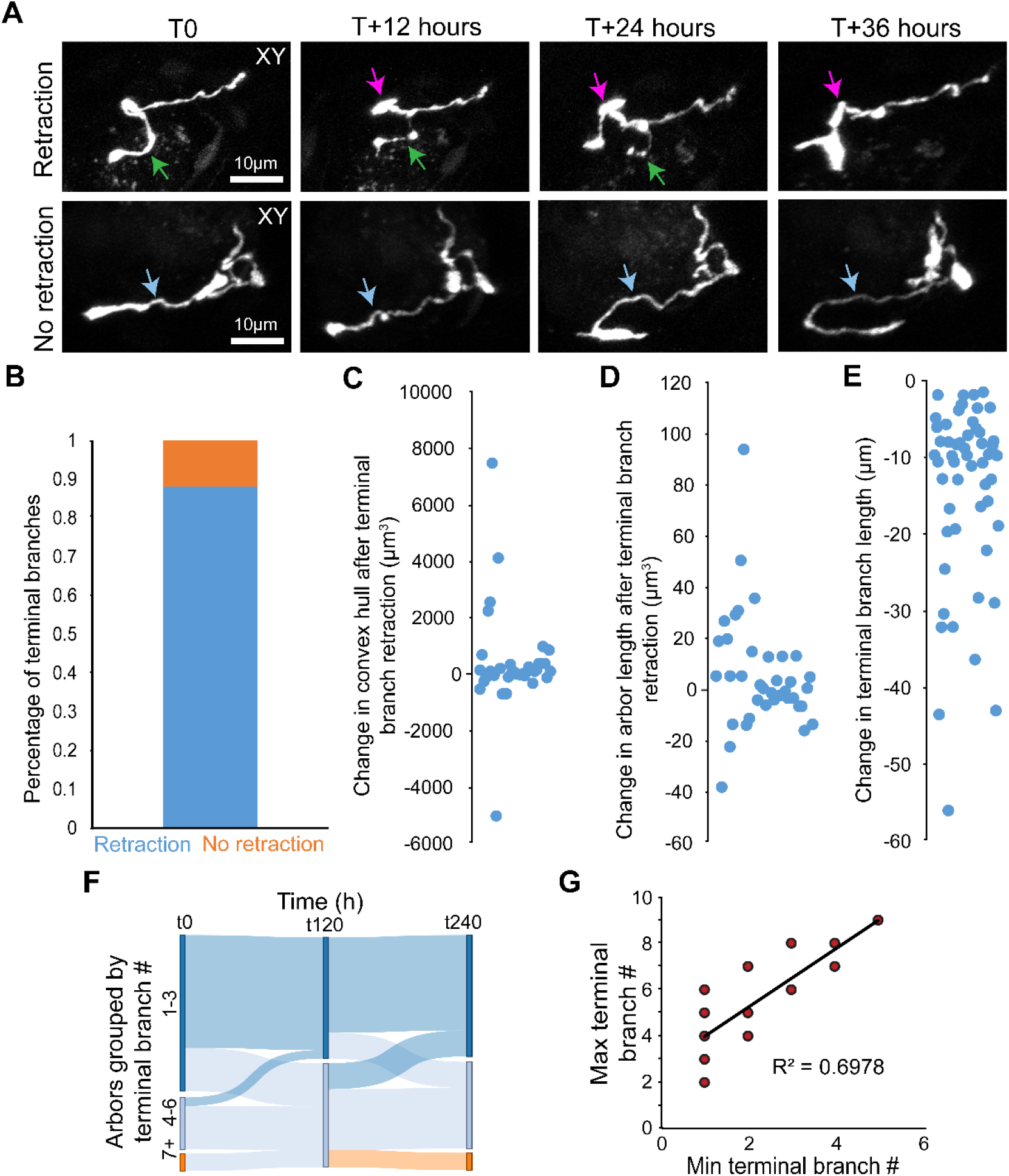
Morphologic changes predict terminal branch retraction but not change in arbor size. (**A**) Two example arbors shown across a 36 h window. In the top row, a terminal branch retracted after displaying a bulb at the tip of the ending and swellings within the branch (green arrow). A separate terminal branch on this same arbor extended (magenta arrow) in conjunction with terminal branch retraction. In the bottom row, a terminal branch displayed an end-bulb and swellings but did not retract (blue arrow); instead, it extended, indicating that the bulb may have been a growth cone. (**B**) Most terminal branches that displayed end-bulbs and swellings retracted at later timepoints (blue); however, a small portion did not retract (orange). (**C**) Change in convex hull after each terminal branch retraction. (**D**) Net change in total arbor length after each terminal branch retraction. (**E**) Change in terminal branch length. (**F**) Sankey diagram showing arbors grouped based on total number of terminal branches (1 to 3 = dark blue, 4 to 6 = light blue, 7+ = orange). Most arbors did not change groups at 5 (t120) or 10 (t240) days of imaging, demonstrating some stability in arbor complexity over time. (**G**) Correlation between minimum and maximum number of terminal branches for 26 receptive arbors observed every 12 h for 10 days.

Since not all branch loss is associated with a retraction bulb, one possibility is that only those retractions with a retraction bulb occur when taste cells are lost. To determine if this was the case, we examined the frequency of retractions which were associated with retraction bulbs. Retractions with retraction bulbs occurred at a frequency of once per 40 h and was also faster than the rate predicted by cell turnover. This finding indicates that branches retract for reasons other than loss of taste bud cells.

We next investigated whether retractions associated with retraction bulbs contributed to reduced arbor size by comparing the volume occupied by the arbor (convex hull; Figure 5C) and arbor length (Figure 5D) before and after terminal branch retraction. Retraction of a single terminal branch did not alter total arbor size (Student’s unpaired t-test, t=0.4515, df=74, p=0.65). These lost branches may represent a small percentage of the total branch length. Consistently, the average length of retraction for a single branch identified using end bulbs and beading was 14.1± 11.9 μm (Figure 5E), which was a small portion of the total length of these arbors (mean=79.2 ± 40.8 μm). In addition, branch elongation also commonly occurred in the arbor at another location concurrently with terminal branch retraction. Consistently, 21 of 38 arbors increased in total length while retracting a terminal branch. These opposing changes in branch length maintain arbor size over time.

Given that arbors add and lose terminal branches concurrently to maintain size, we wondered to what extent the total range of arbor complexity, measured as terminal branch number, changed in each arbor over 10 days. We observed that most arbors maintained a limited range of complexity over 10 days (Figure 5F), such that the minimum and maximum number of terminal branches for each arbor was correlated (Figure 5G, R^2^ = 0.698, p < 0.001). Simple arbors (1-3 branches) did not become complex (more than 6 branches) within a 10-day window. Although large branch retractions also occurred frequently within 10 days, they occurred simultaneously with terminal branch formation, which limited the range of complexity for most arbors.

### Variability in arbor size over 10 days differs across arbors

Because arbor size (convex hull) was not heavily impacted by the retraction of single terminal branches we hypothesized that arbor size is maintained despite constant arbor remodeling. To determine if this was the case, we examined whether arbor size (convex hull and total arbor length) changed over 10 days. To perform comparisons across all timepoints within a 10-day period, we developed a semi-automated image analysis pipeline in MATLAB, called ArborTools, to segment arbors and quantify convex hull and total arbor length (Figure 6A). Reconstructions of each arbor were verified visually to confirm that the arbor and the reconstruction matched (Figure 6, cyan vs white). Arbors with too much background noise for automatic tracing were removed from analysis. The volume occupied by the arbor (convex hull) changed in the taste bud across 10 days. In general, convex hull changed along with total length (Figure 6C). These changes in size could be due to changes in height, or x or y width, or combinations of these changes (Figure 6D). For example, the illustrated arbor (Figure 6 A-C), increases in size occur due to an increase in Y-width from 0 to 12 hours, but dramatically increases in size between hour 228-240 due to concurrent increases X, Y width and height.

**Figure 6.**
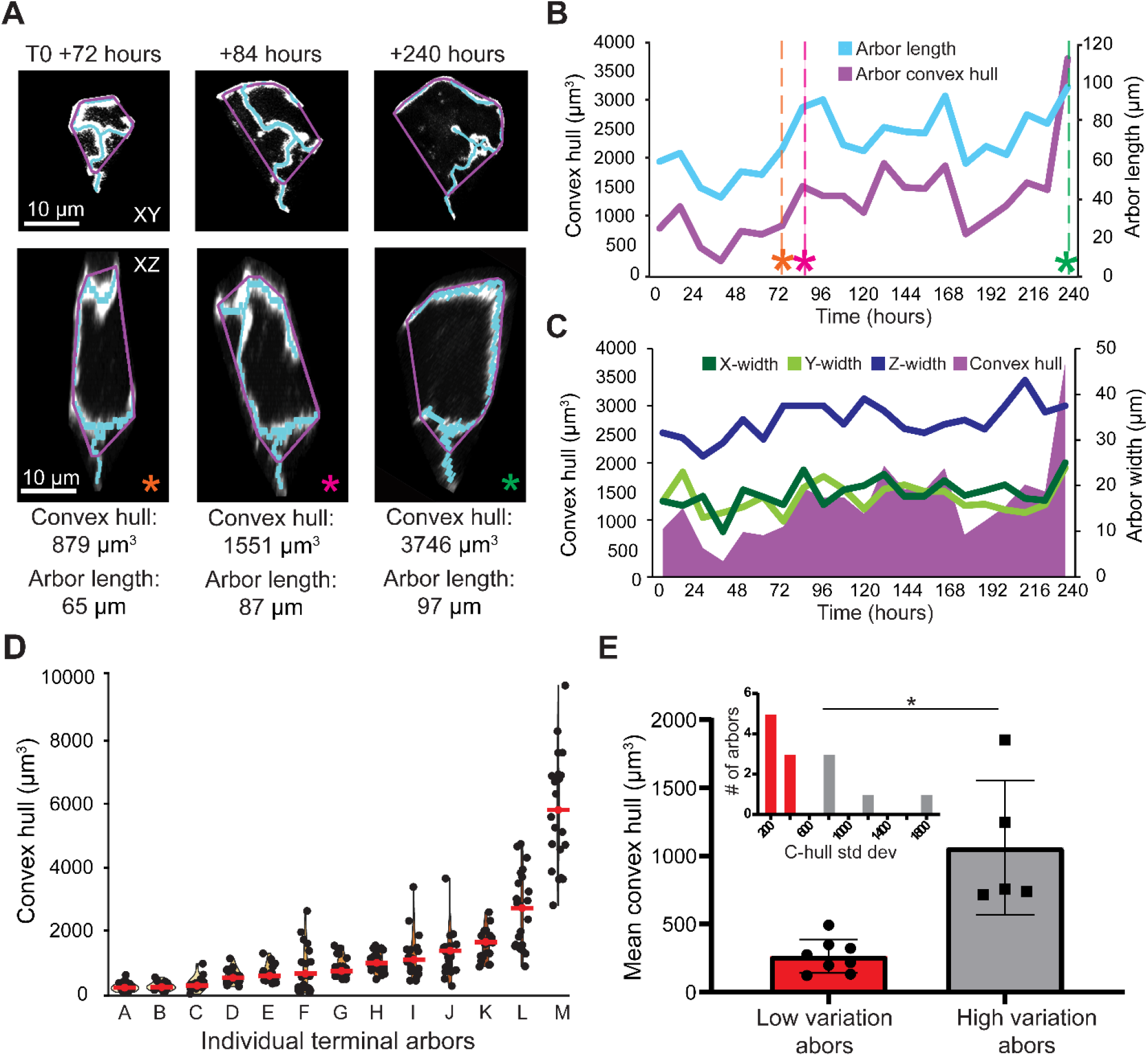
Taste neuron arbors vary in size over 10 days. A custom image analysis pipeline was developed to segment arbor structure from raw image TIF stacks. (**A**) Examples of convex hulls (purple) and arbor skeletons (cyan) generated from segmentation data are shown in two orientations and at three timepoints. Colored asterisks at each timepoint are used to indicate convex hull and arbor length in (C). (**B**) Convex hull size (purple line) and arbor length (cyan) for the example arbor in (A) are shown across the entire 10-day imaging window. **(C)** Convex hull potted with Z-height, x-width and y-width over time. (**D**) Convex hull measured for 13 individual arbors arranged by ascending average size. Average convex hull and arbor length during the 10-day imaging window are indicated with red dots and bars. **(E)** Within 10 days small arbors remodel little and are small, while other remodel more extensively and tend to be larger and more variant in size.

When examining the extent to which individual arbors displayed changes in size, we noticed that some arbors appear to be more variable (Figure 6D). We divided them into two groups based on the variability of their sizes over 10 days (Figure 6E, insert). Arbors appear to divide into those that vary little over time and remain small, and those that remodel more extensively (Figure 6E). Those that remodel more extensively are also larger on average over 10 days. These two groups could represent phases in a remodeling cycle that extends beyond 10 days, or it could represent two distinct populations of taste arbors that differ is regard to degree of remodeling.

### Number of arbors within a taste bud is stable over 100 days

Within 10-days, we did not observe any new arbors growing into taste buds, and only a small number of arbors were lost. Therefore, total number of arbors may be a stable feature for a given taste ganglion neuron over time. Alternatively, gain loss of whole arbors may occur, but require a longer time than 10 days. To distinguish between these possibilities, we imaged taste buds every 10 days for 60 or 100 days in mice where neurons were sparsely labeled. We observed taste buds with labeled arbors (n = 18 for 60 days, n = 13 for 100 days) as well as buds without labeled arbors (n = 18 for 60 days, n = 13 for 100 days) (Figure 7A). The arbor structure in the papillae core was largely unchanged over 100 days (Figure 7B, purple dotted line). In addition, no taste buds acquired new labeled arbors in 60 or 100 days of imaging. Initially, 18 taste buds contained a total of 33 labeled arbors. After 60 days of imaging, 31 arbors were still present. No additional labeled arbors were lost from the remaining taste buds throughout the 100 days (Figure 7C). These results suggest that the number of arbors is a stable feature of the neuron in adulthood. Arbors do not exhibit the full range of complexity over 100 days (Figure 7D). Convex hull and arbor length also indicate that some arbors remain small, while others are more variable (Figure 7E-F).

**Figure 7.**
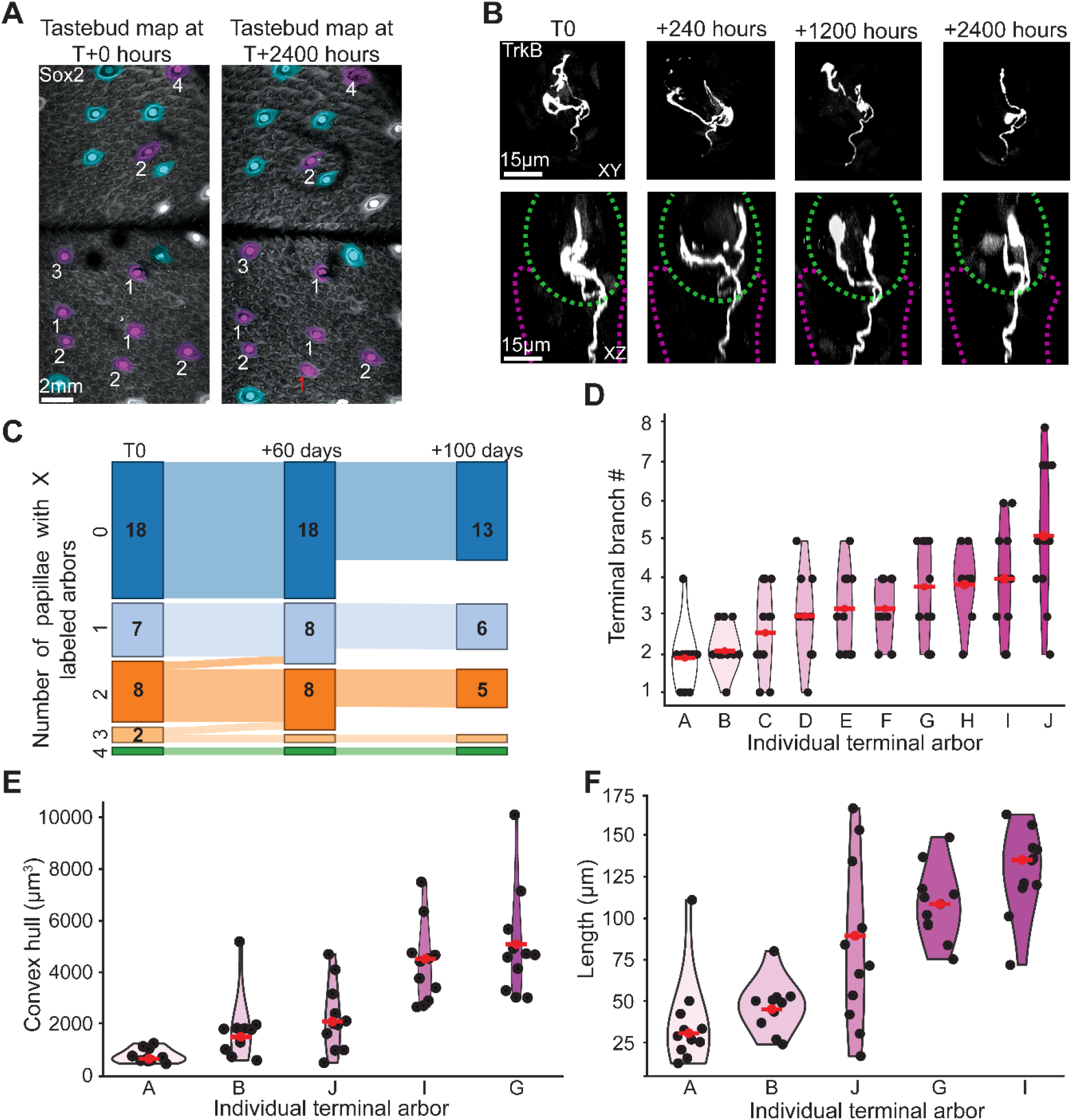
Arbor number is stable over 100 days. (**A**) Two papillae maps of the same regions shown 100 days apart. Purple taste buds contained labeled arbors (number of arbors labeled in white), and blue taste buds did not contain labeled arbors. One papilla in this map lost an arbor during the 100-day window (red label). (**B**) Example arbor at days 0, 10, 50, and 100. The taste bud is outlined in a green dotted line and papillae core in magenta. (**C**) Taste buds were categorized by number of labeled arbors and plotted in a Sankey diagram. No taste buds acquired new arbors, and two taste buds lost an arbor during the first 60 days. One mouse was not imaged past day 60. (**D**) Violin plots of terminal branch number for 10 arbors imaged every 10 days for 100 days. Average number of terminal branches during the 100-day window are indicated with red dots and bars. Violin plots of convex hull (**E**) and arbor length (**F**) for five individual arbors imaged every 10 days for 100 days. Arbor identities are conserved from (**D**).

## Discussion

Taste bud cells are continuously replaced through adulthood (Beidler and Smallman, 1965; Farbman, 1980; Hamamichi et al., 2006), but whether taste ganglion neurons remodel while reconnecting with new taste bud cells was unknown. In other systems, intra-vital imaging has been used to examine neuron remodeling during development, following injury, and in some cases during adulthood (Breton-Provencher et al., 2016; De Paola et al., 2006; Marshall et al., 2016; Nakazawa et al., 2018; Pfeiffer et al., 2018; Speidel, 1933, 1932; Takahashi et al., 2019; Trachtenberg et al., 2002; Verzé et al., 1999; Walsh and Lichtman, 2003). Here we show that all peripheral taste axons have arbors that continuously remodel, and compared to plasticity in other systems, do so with unprecedented speed. For example, a structure well-studied for its remodeling capabilities in adulthood, the dendritic spine, remodels over days (Pfeiffer et al., 2018), and at least half of the observed spines are stable over months (Grutzendler et al., 2002; Trachtenberg et al., 2002). In contrast to dendritic spines, some terminal branches of taste axonal arbors appear and leave over much shorter time-scales – sometimes lasting less than 12 hours. The speed and extent of remodeling in peripheral taste axons provides insight into two basic questions. First, how do nerve arbors locate a new taste bud cell during the process taste cell renewal? Secondly, is morphological variation across individual peripheral axons due entirely to plasticity, or can taste neurons be classified as distinct types based on anatomical differences? We discuss each of these issues.

### The relationship between taste arbor remodeling and cell renewal

Most taste arbors only connect to one taste bud cell at any given time (Kinnamon et al., 1988; Wilson et al., 2022). In a 10-day period roughly half of the population of taste bud cells turnover. Therefore, we anticipated that approximately half of all arbors sampled would lose their connection to a bud cell and/or gain a new connection within 10-days. We expected that arbor structure would be stable while connected to a taste bud cell, remodeling only when connections were broken (Ohman et al., submitted). This type of phasic remodeling is observed in terminal arbors of touch neurons that receive input from Merkel cells (Marshall et al., 2016). The results of our study do not support the model that taste neurons experience distinct phases of remodeling and stability; rather the arbors of taste ganglion neurons continuously remodel with concurrent terminal branch growth and retraction, similar to the pattern of remodeling in touch terminals of ear-skin (Takahashi et al., 2019).

During remodeling, the arbors of taste ganglion neurons display similar anatomical features to motor axon terminals, which innervate neuromuscular junctions, and undergo remodeling as refinement occurs during development. However, refinement largely consists of terminal branch retraction and pruning (Bishop et al., 2004; Walsh and Lichtman, 2003). Motor axon retraction is typically accompanied by retraction bulbs and terminal branch swellings. In addition to refinement, these features are also seen during degeneration after injury (Bernstein and Lichtman, 1999; Bishop et al., 2004; Erturk et al., 2007). Similarly, the axonal arbors of taste ganglion neurons have also been described as having these irregular swellings and end-bulbs (Zaidi et al., 2016). Here, we found that branches displaying these features are indeed in the process of retracting. These observations suggest that retraction occurs by a conserved mechanism across systems – a process that involves axosomal vesicle shedding and engulfment of neuronal debris by phagocytic cells (Bishop et al., 2004). In the case of motor neurons, there is evidence that factors provided from the neuromuscular junction trigger terminal branch retraction (Bernstein and Lichtman, 1999). While the anticipated source of such a signal in the taste bud would be from a dying taste bud cell, retraction occurs more frequently than would be predicted based on the rate at which cells are leaving the taste bud. The frequency of terminal branch retractions suggests that a signal from a dying taste bud cell is not required to trigger terminal branch retraction in taste neuron terminal arbors.

Taste bud cells release growth and guidance factors, and as new cells enter the taste bud they could release factors that stimulate branch formation and/or extension (Donnelly et al., 2018; Lee et al., 2017; Meng et al., 2015; Nosrat et al., 2012; Sukumaran et al., 2017; Treffy et al., 2016; Yee et al., 2003). However, our finding that terminal branch remodeling is much faster than expected based on the rate of cell turnover is not consistent with this possibility. Instead, terminal branch remodeling may be an intrinsic feature of taste ganglion neurons, occurring regardless of what external factors are present. To assess whether this remodeling is driven by an intrinsic mechanism, we examined remodeling of terminal arbors while preventing addition of new cells into taste buds. Treatment with LDE225 inhibits taste bud cell differentiation such that new taste bud cells do not enter the taste bud (Castillo-Azofeifa et al., 2017; Kumari et al., 2015). Surprisingly, we found that preventing integration of new taste bud cells does not influence the frequency of terminal branch loss or formation. In fact, we observed that taste arbors remain and continue to remodel even after the entire taste bud was lost. This finding is consistent with the presence of growth associated protein-43 (GAP43) in the arbors following treatment with LDE225 (Donnelly et al., 2022). Collectively, these findings support the conclusion that axonal arbor remodeling is an intrinsic characteristic of peripheral taste neurons.

Previously, we hypothesized taste arbors remodel to connect with new taste cells over time (Ohman and Krimm, 2021). This model assumed that taste bud cells provide an extrinsic signal to influence arbor remodeling. Because this concept is not supported experimentally by the current study, we offer an updated model of how taste nerve fibers connect to new taste bud cells over time. We propose that axonal arbors of taste ganglion neurons continuously sample the taste bud compartment by extending and retracting terminal branches, and this remodeling is orchestrated by an intrinsic cellular mechanism. An intrinsic program that drives constant remodeling allows taste arbors to detect extracellular signals and compare spatial arrangements of taste cells in their immediate environment. By extending branches randomly and continuously nerve arbors eventually come near to a cell of the “correct type”. Once it is in the vicinity of the new taste bud cell, trophic mechanisms may attract the taste arbor a short distance to the cell. Furthermore, if constant terminal branch remodeling causes a terminal arbor to come in contact with a cell, then detection of surface molecules, such as protocadherin could regulate the connection, and growth factors may be unnecessary (Hirose et al., 2020). In this scenario, terminal branch remodeling orchestrates the detection of extrinsic molecular cues and is not a response by peripheral taste neurons to extrinsic molecular cues.

### The role of plasticity in individual peripheral taste axon morphology

A primary goal of the current study was to determine which structural features of peripheral taste neurons are plastic and which features are stable. Addressing this question is fundamental in determining whether taste neurons can be classified morphologically into separate types (Dvoryanchikov et al., 2017; Zhang et al., 2019). Taste neurons are pseudo-unipolar sensory neurons with a single axon and a primary bifurcation that creates a central and peripheral projection (Yarmolinsky et al., 2009; Zaidi and Whitehead, 2006). We recently described the morphology of taste neurons for the first time using a fixed-tissue approach (Huang et al., 2021). The peripheral axons of taste neurons vary considerably in structure due to differences in the number of branch points within the tongue muscle, the lamina propria and within the taste bud (Huang et al., 2021). Here we show that the taste arbors remodel, and the number terminal branches for an individual axonal arbor is determined by plasticity.

Although terminal branches remodel quickly, the gain and loss of terminal branches occur concurrently, such that loss of a terminal branch does not always reflect a decrease in arbor size. Individual arbor size does vary over time; however, some individual arbors change in size more than others. We recently found that individual taste neurons either have all small arbors, or have a mix of large and small arbors (Ohman et al., in submission). It may be that degree of plasticity in an arbor size varies across different types of taste neurons. Large arbors with higher degrees of plasticity likely probe more of the taste bud environment, and are more likely to locate a scarce taste bud cell type. So these type of endings could be specialized to neuron types that are narrowly-tuned and require input from uncommon taste cell types.

Interestingly, the number of arbors labeled remained stable over 100 days. The presence or absence of arbors in the bud is dictated by branch points within the axon that are in the muscle or lamina propria. These branch points are not added and/or lost during adulthood. Even when taste buds are eliminated, individual arbors remain in the location previously occupied by taste buds. The axons of taste ganglion neurons may have between 1 to 14 arbors (Huang et al., 2021). Given that most arbors only connect to 1-2 cells, the number of arbors is the primary anatomical feature that determines how many taste bud cells provide input to a given neuron (Wilson et al., 2022). Neurons with more arbors are predicted to be both more sensitive and broadly tuned than neurons with fewer arbors (Ohman and Krimm, 2021). Although there is evidence that peripheral taste neurons can change function over time (Bradley et al., 1997), there is also likely some level of functional stability so that taste information is relayed to the brain as an interpretable code. Maintaining a constant number of arbors is one mechanism by which taste neurons could maintain functional stability over time.

Here, we determined that some anatomical features of peripheral taste axons are highly plastic while others display long-term stability. Stable anatomical features of taste ganglion neurons that also vary across neurons could define morphological neuron types. These stable morphological differences may correspond to genetically defined neuron types (Dvoryanchikov et al., 2017; Zhang et al., 2019) or functional differences (Barretto et al., 2015; Lundy and Contreras, 1999; Wu et al., 2015). However, It is still unclear whether peripheral taste neuron types are defined by a combination of expression, function, and stable morphological features (Zeng and Sanes, 2017). By describing which portions of taste axons are plastic and which are stable, we are now poised to determine if stable morphological features differ across genetic or functional types.

## Materials and methods

### Animals

All animals were cared for in accordance with guidelines set by the United States Public Health Service Policy on the Humane Care and Use of Laboratory Animals and the National Institutes of Health Guide for the Care and Use of Laboratory Animals.

*TrkB*^CreER^ mice (*Ntrk2*^*tm3*.*1(cre/ERT2)Ddg*^; ; ISMR catalog #JAX:027214, RRID:IMSR_JAX:027214) were crossed with Cre-dependent tdTomato mice (Rutlin et al., 2014; RRID: IMSR_JAX:007914) to obtain *TrkB*^CreER^:tdTomato mice in which tdTomato is expressed following TrkB-driven Cre-mediated gene recombination. These mice were crossed with a Sox2-GFP line (Sox2^tm2Hoch^;; ISMR catalog #JAX:017592, RRID:IMSR_JAX:017592). Phox2b-Cre mice (MRRC_034613-UCD) were also bred with tdTomato reporter mice for fixed tissue experiments. K14^CreERT^ mice; ISMR catalog #JAX:005107, RRID:IMSR_JAX:005107) were crossed with tdTomato reporter mice for lineage tracing of taste cells. For all experiments, mice were postnatal day 60 or older at the initial imaging session. Males and females were used across all *in vivo* experiments (24 males and 19 females in total). Mice were divided across experiments as follows: 10-day imaging (n = 5), 100-day imaging (n = 3). 4 h imaging (n = 5) 1 h imaging (n = 4), 6-h imaging (n = 2), 4 h live imaging Hh-inhibition (n = 6; vehicle n = 2, LDE225 n = 4), new cells entering the taste bud (n = 7; vehicle n = 2, LDE225 n = 5), confocal imaging experiments examining taste bud size (n = 10; vehicle n = 5, LDE225 n = 5).

### Tamoxifen administration

Tamoxifen (T5648, MiliporeSigma, St. Louis, MO) was dissolved in corn oil (C-8267, MiliporeSigma) at 20 mg/ml by shaking and heating at 42°C and injected in a single dose on postnatal day 40 by intragastric gavage.

### LDE225 administration

Mice were treated by daily oral gavage for 10, 16, or 23 days with vehicle (PEG 400:5% dextrose in water) or LDE225 (NVP-LDE225, ChemieTek, Indianapolis, IN) dissolved in vehicle at a dose of 20 mg/kg. The dose was determined based on Kumari et al. (2015) and preliminary studies. The oral gavage probe (22-gauge, 25 mm; Instech) by-passed all oral tissues for direct delivery into the stomach.

### Immunohistochemistry

Phox2b-Cre:tdTomato mice were killed by avertin overdose (4 mg/kg) and perfused transcardially with 4% paraformaldehyde. Dissected tissues were postfixed in 4% paraformaldehyde for 2 h (for thin serial sections) or overnight (thick sections and whole mounts), rinsed with PBS, and transferred to 30% sucrose at 4°C overnight. A razor blade was used to remove the anterior two-thirds of the tongue, after which tongues were carefully split down the midline with a razor blade under a dissection microscope. Tongues were frozen the next day in OCT and stored at −80°C before processing for whole-mount staining. Whole-mount immunohistochemistry of the lingual epithelium was performed to visualize innervated taste buds. First, the underlying muscle and lamina propria were removed as described previously (Ohman and Krimm, 2021). The isolated lingual epithelium was then washed for 15 min (three times) in 0.1 M PBS. Tissues were then incubated in blocking solution (3% donkey serum and 0.5% Triton X-100 in 0.1 M PBS) at 4°C overnight and then incubated for 5 days at 4°C with primary antibodies (DsRed (rabbit) and Troma1 (rat)) in antibody solution (0.5% Triton X-100 in 0.1 M PBS). Tissues were rinsed four times for 15 min each with 0.1 M PBS and incubated with secondary antibodies anti-DsRed (1:500; RRID:AB_10013483; Living Colors DsRed polyclonal; Takara Bio USA, San Jose, CA) and anti-Troma1 (1:500, Alexa Fluor 488 AffiniPure, RRID:AB_2340619; Jackson ImmunoResearch, West Grove, PA). Tissues were then rinsed again, mounted with Fluoromount-G, and coverslipped (high precision, 0107242; Marienfeld).

### Confocal imaging

Taste bud images were obtained using an Olympus Fluoview FV1000 confocal laser-scanning microscope with a 60× NA1.4 lens objective using a zoom of 3, Kalman 2. Image sizes were initially set at 1024 × 1024 pixels but were cropped to reduce scanning time and bleaching. Serial optical sections at intervals of 0.47 μm in the Z dimension were captured, which is the optimal size at 60× magnification for 3D reconstruction. All colors were imaged sequentially in separate channels to avoid bleed-through. Image stacks were then deconvolved using AutoQuant X3 software (Media Cybernetics, Rockville, MD) to reduce out-of-focus fluorescence and improve image quality.

### Two-photon *in vivo* imaging

*In vivo* 2PLSM imaging was performed using a movable objective microscope (Sutter Instruments, Novato, CA). A Fidelity-2 1070-nm laser (Coherent, Silicon Valley was used to visualize tdTomato, and a Chameleon tunable laser (Coherent) set to 920 nm was used to visualize GFP. A custom-built tongue holder, based on a published design, was used to stabilize the anterior tongue for imaging of the dorsal surface (Han and Choi, 2018). Mice were anesthetized using a 0.6% isoflurane (Henry Schien, Melville, NY) and O_2_ mixture. A temperature controller was used to monitor and maintain body temperature at 36°C. Ophthalmic lubricant ointment (Henry Schien) was applied to the eyes. Taste bud maps were acquired using a 10 × 0.3 numerical aperture water immersion objective lens (Carl Zeiss, USA), and images of receptive arbors were acquired using a 40 × 10 numerical aperture water immersion lens (Zeiss). An early version of ScanImage (Matlab) was used to collect images. Receptive arbors were typically imaged to 70 microns in depth below the surface of the tongue, and 80 × 80 μm optical sections (512 × 512 pixels) were collected at 0.5 or 1 μm increments.

### Image analysis

Taste bud volume was calculated by contouring whole-mount-stained taste buds at 3 micron increments perpendicular to the XY plane of the confocal z-stack. K14 cells were manually quantified from taste bud z-stacks using ImageJ. Terminal branches of sparsely labeled receptive arbors were manually quantified from z-stacks using ImageJ. Terminal branches that displayed retraction bulbs were quantified using the SNT plugin. A custom Matlab program (ArborTools, Github link to software will go here) was used to automatically segment arbors. 3D images stacks were cropped to isolate terminal arbors from background. A custom filter was used to extract initial segmentation volumes (Lang et al., 2011). After automatic segmentation, the segmented arbors were compared to the arbors and instances where background pixels were segmented were manually selected to not be included in the convex hull and length analysis. From segmented volumes skeletons were extracted using the Treestoolbox (Cuntz et al., 2010). All convex hull and arbor length data reported were generated from ArborTools.

### Statistical analysis

Rates of terminal branch change were compared across image sets by first testing whether the measure was normally distributed using a Shapiro-Wilk test. If a measure was normally distributed, then differences were determined using a Student’s t-test for two groups or one-way analysis of variance with a Tukey *post hoc* test for more than two groups. If a measure was not normally distributed, then differences were determined using a Mann-Whitney test for two groups or Kruskal-Wallis test for more than two groups. A p-value <0.05 was used to determine statistical significance.

## Supplementary material

S1 Movie. The arbor from Figure 3A (x-Y view) from T0-T228, every 12 hours = 0.5 seconds Time in hours is on the bottom right-hand side of the images. Branches are added and lost to the portion of the arbor inside the taste bud.

## References

AU-Ohman LC, AU-Krimm RF. 2021. Whole-Mount Staining, Visualization, and Analysis of Fungiform, Circumvallate, and Palate Taste Buds. J Vis Exp e62126. doi:10.3791/62126

Barretto RPJ, Gillis-Smith S, Chandrashekar J, Yarmolinsky DA, Schnitzer MJ, Ryba NJP, Zuker CS. 2015. The neural representation of taste quality at the periphery. Nature 517:373–376. doi:10.1038/nature13873

Beidler LM, Smallman RL. 1965. RENEWAL OF CELLS WITHIN TASTE BUDS. J Cell Biol 27:263–272. doi:10.1083/jcb.27.2.263

Bernstein M, Lichtman JW. 1999. Axonal atrophy: The retraction reaction. Curr Opin Neurobiol 9:364–370. doi:10.1016/S0959-4388(99)80053-1

Bishop DL, Misgeld T, Walsh MK, Gan W-B, Lichtman JW. 2004. Axon Branch Removal at Developing Synapses by Axosome Shedding. Neuron 44:651–661. doi:10.1016/j.neuron.2004.10.026

Bradley RM, Cao X, Akin T, Najafi K. 1997. Long term chronic recordings from peripheral sensory fibers using a sieve electrode array. J Neurosci Methods 73:177–186. doi:10.1016/S0165-0270(97)02225-5

Breton-Provencher V, Bakhshetyan K, Hardy D, Bammann RR, Cavarretta F, Snapyan M, Côté D, Migliore M, Saghatelyan A. 2016. Principal cell activity induces spine relocation of adult-born interneurons in the olfactory bulb. Nat Commun 7:12659. doi:10.1038/ncomms12659

Castillo D, Seidel K, Salcedo E, Ahn C, de Sauvage FJ, Klein OD, Barlow LA. 2014. Induction of ectopic taste buds by SHH reveals the competency and plasticity of adult lingual epithelium. Development 141:2993–3002. doi:10.1242/dev.107631

Castillo-Azofeifa D, Losacco JT, Salcedo E, Golden EJ, Finger TE, Barlow LA. 2017. Sonic hedgehog from both nerves and epithelium is a key trophic factor for taste bud maintenance. Development 144:3054–3065. doi:10.1242/dev.150342

Cuntz H, Forstner F, Borst A, Häusser M. 2010. One Rule to Grow Them All: A General Theory of Neuronal Branching and Its Practical Application. PLoS Comput Biol 6:e1000877. doi:10.1371/journal.pcbi.1000877

De Paola V, Holtmaat A, Knott G, Song S, Wilbrecht L, Caroni P, Svoboda K. 2006. Cell Type-Specific Structural Plasticity of Axonal Branches and Boutons in the Adult Neocortex. Neuron 49:861–875. doi:10.1016/j.neuron.2006.02.017

Donnelly CR, Kumari A, Li L, Vesela I, Bradley RM, Mistretta CM, Pierchala BA. 2022. Probing the multimodal fungiform papilla: complex peripheral nerve endings of chorda tympani taste and mechanosensitive fibers before and after Hedgehog pathway inhibition. Cell Tissue Res 387:225–247. doi:10.1007/s00441-021-03561-1

Donnelly CR, Shah AA, Mistretta CM, Bradley RM, Pierchala BA. 2018. Biphasic functions for the GDNF-Ret signaling pathway in chemosensory neuron development and diversification. Proc Natl Acad Sci 115. doi:10.1073/pnas.1708838115

Dvoryanchikov G, Hernandez D, Roebber JK, Hill DL, Roper SD, Chaudhari N. 2017. Transcriptomes and neurotransmitter profiles of classes of gustatory and somatosensory neurons in the geniculate ganglion. Nat Commun 8. doi:10.1038/s41467-017-01095-1

Ermilov AN, Kumari A, Li L, Joiner AM, Grachtchouk MA, Allen BL, Dlugosz AA, Mistretta CM. 2016. Maintenance of Taste Organs Is Strictly Dependent on Epithelial Hedgehog/GLI Signaling. PLOS Genet 12:e1006442. doi:10.1371/journal.pgen.1006442

Erturk A, Hellal F, Enes J, Bradke F. 2007. Disorganized Microtubules Underlie the Formation of Retraction Bulbs and the Failure of Axonal Regeneration. J Neurosci 27:9169–9180. doi:10.1523/JNEUROSCI.0612-07.2007

Farbman AI. 1980. RENEWAL OF TASTE BUD CELLS IN RAT CIRCUMVALLATE PAPILLAE. Cell Prolif 13:349–357. doi:10.1111/j.1365-2184.1980.tb00474.x

Gangadharan V, Zheng H, Taberner FJ, Landry J, Nees TA, Pistolic J, Agarwal N, Männich D, Benes V, Helmstaedter M, Ommer B, Lechner SG, Kuner T, Kuner R. 2022. Neuropathic pain caused by miswiring and abnormal end organ targeting. Nature 606:137–145. doi:10.1038/s41586-022-04777-z

Grutzendler J, Kasthuri N, Gan W-B. 2002. Long-term dendritic spine stability in the adult cortex. Nature 420:812–816. doi:10.1038/nature01276

Hamamichi R, Asano-Miyoshi M, Emori Y. 2006. Taste bud contains both short-lived and long-lived cell populations. Neuroscience 141:2129–2138. doi:10.1016/j.neuroscience.2006.05.061

Han J, Choi M. 2018. Comprehensive functional screening of taste sensation in vivo (preprint). Neuroscience. doi:10.1101/371682

Hirose F, Takai S, Takahashi I, Shigemura N. 2020. Expression of protocadherin-20 in mouse taste buds. Sci Rep 10:2051. doi:10.1038/s41598-020-58991-8

Huang T, Ohman LC, Clements AV, Whiddon ZD, Krimm RF. 2021. Variable Branching Characteristics of Peripheral Taste Neurons Indicates Differential Convergence. J Neurosci 41:4850. doi:10.1523/JNEUROSCI.1935-20.2021

Jacobsen AA, Aldahan AS, Hughes OB, Shah VV, Strasswimmer J. 2016. Hedgehog Pathway Inhibitor Therapy for Locally Advanced and Metastatic Basal Cell Carcinoma: A Systematic Review and Pooled Analysis of Interventional Studies. JAMA Dermatol 152:816. doi:10.1001/jamadermatol.2016.0780

Kinnamon JC, Sherman TA, Roper SD. 1988. Ultrastructure of mouse vallate taste buds: III. Patterns of synaptic connectivity. J Comp Neurol 270:1–10. doi:10.1002/cne.902700102

Kumari A, Ermilov AN, Allen BL, Bradley RM, Dlugosz AA, Mistretta CM. 2015. Hedgehog pathway blockade with the cancer drug LDE225 disrupts taste organs and taste sensation. J Neurophysiol 113:1034–1040. doi:10.1152/jn.00822.2014

Kumari A, Ermilov AN, Grachtchouk M, Dlugosz AA, Allen BL, Bradley RM, Mistretta CM. 2017. Recovery of taste organs and sensory function after severe loss from Hedgehog/Smoothened inhibition with cancer drug sonidegib. Proc Natl Acad Sci 114:E10369–E10378. doi:10.1073/pnas.1712881114

Lacouture ME, Dréno B, Ascierto PA, Dummer R, Basset-Seguin N, Fife K, Ernst S, Licitra L, Neves RI, Peris K, Puig S, Sokolof J, Sekulic A, Hauschild A, Kunstfeld R. 2016. Characterization and Management of Hedgehog Pathway Inhibitor-Related Adverse Events in Patients With Advanced Basal Cell Carcinoma. The Oncologist 21:1218–1229. doi:10.1634/theoncologist.2016-0186

Lang S, Drouvelis P, Tafaj E, Bastian P, Sakmann B. 2011. Fast extraction of neuron morphologies from large-scale SBFSEM image stacks. J Comput Neurosci 31:533–545. doi:10.1007/s10827-011-0316-1

Lee H, Macpherson LJ, Parada CA, Zuker CS, Ryba NJP. 2017. Rewiring the taste system. Nature 548:330–333. doi:10.1038/nature23299

Liebl DJ, Mbiene J-P, Parada LF. 1999. NT4/5 Mutant Mice Have Deficiency in Gustatory Papillae and Taste Bud Formation. Dev Biol 213:378–389. doi:10.1006/dbio.1999.9385

Lindemann B. 2001. Receptors and transduction in taste. Nature 413:219–225. doi:10.1038/35093032

Lu W-J, Mann RK, Nguyen A, Bi T, Silverstein M, Tang JY, Chen X, Beachy PA. 2018. Neuronal delivery of Hedgehog directs spatial patterning of taste organ regeneration. Proc Natl Acad Sci 115:E200–E209. doi:10.1073/pnas.1719109115

Lundy RF, Contreras RJ. 1999. Gustatory Neuron Types in Rat Geniculate Ganglion. J Neurophysiol 82:2970–2988. doi:10.1152/jn.1999.82.6.2970

Marshall KL, Clary RC, Baba Y, Orlowsky RL, Gerling GJ, Lumpkin EA. 2016. Touch Receptors Undergo Rapid Remodeling in Healthy Skin. Cell Rep 17:1719–1727. doi:10.1016/j.celrep.2016.10.034

Martineau É, Di Polo A, Vande Velde C, Robitaille R. 2018. Dynamic neuromuscular remodeling precedes motor-unit loss in a mouse model of ALS. eLife 7:e41973. doi:10.7554/eLife.41973

Meng L, Ohman-Gault L, Ma L, Krimm RF. 2015. Taste Bud-Derived BDNF Is Required to Maintain Normal Amounts of Innervation to Adult Taste Buds. eneuro 2:ENEURO.0097-15.2015. doi:10.1523/ENEURO.0097-15.2015

Miller IJ, Reedy FE. 1990. Variations in human taste bud density and taste intensity perception. Physiol Behav 47:1213–1219. doi:10.1016/0031-9384(90)90374-D

Nakazawa S, Mizuno H, Iwasato T. 2018. Differential dynamics of cortical neuron dendritic trees revealed by long-term in vivo imaging in neonates. Nat Commun 9:3106. doi:10.1038/s41467-018-05563-0

Nosrat IV, Margolskee RF, Nosrat CA. 2012. Targeted Taste Cell-specific Overexpression of Brain-derived Neurotrophic Factor in Adult Taste Buds Elevates Phosphorylated TrkB Protein Levels in Taste Cells, Increases Taste Bud Size, and Promotes Gustatory Innervation. J Biol Chem 287:16791–16800. doi:10.1074/jbc.M111.328476

Ohman LC, Hanbala L, Krimm RF. in submission. Taste arbor structural variability analyzed across taste regions.

Ohman LC, Krimm RF. 2021. Variation in taste ganglion neuron morphology: insights into taste function and plasticity. Curr Opin Physiol 20:134–139. doi:10.1016/j.cophys.2020.12.011

Ohman-Gault L, Huang T, Krimm R. 2017. The transcription factor Phox2b distinguishes between oral and non-oral sensory neurons in the geniculate ganglion. J Comp Neurol 525:3935–3950. doi:10.1002/cne.24312

Okubo T, Pevny LH, Hogan BLM. 2006. Sox2 is required for development of taste bud sensory cells. Genes Dev 20:2654–2659. doi:10.1101/gad.1457106

Patel AV, Krimm RF. 2010. BDNF is required for the survival of differentiated geniculate ganglion neurons. Dev Biol 340:419–429. doi:10.1016/j.ydbio.2010.01.024

Pfeiffer T, Poll S, Bancelin S, Angibaud J, Inavalli VK, Keppler K, Mittag M, Fuhrmann M, Nägerl UV. 2018. Chronic 2P-STED imaging reveals high turnover of dendritic spines in the hippocampus in vivo. eLife 7:e34700. doi:10.7554/eLife.34700

Purves D, Hadley R. 1985. Changes in the dendritic branching of adult mammalian neurones revealed by repeated imaging in situ. Nat Lond 315:404–406.

Romanov RA, Lasher RS, High B, Savidge LE, Lawson A, Rogachevskaja OA, Zhao H, Rogachevsky VV, Bystrova MF, Churbanov GD, Adameyko I, Harkany T, Yang R, Kidd GJ, Marambaud P, Kinnamon JC, Kolesnikov SS, Finger TE. 2018. Chemical synapses without synaptic vesicles: Purinergic neurotransmission through a CALHM1 channel-mitochondrial signaling complex. Sci Signal 11:eaao1815. doi:10.1126/scisignal.aao1815

Speidel CC. 1935. STUDIES OF LIVING NERVES: IV. GROWTH, REGENERATION, AND MYELINATION OF PERIPHERAL NERVES IN SALAMANDERS. Biol Bull 68:140–161. doi:10.2307/1537291

Speidel CC. 1933. Studies of living nerves. II. Activities of ameboid growth cones, sheath cells, and myelin segments, as revealed by prolonged observation of individual nerve fibers in frog tadpoles. Am J Anat 52:1–79. doi:10.1002/aja.1000520102

Speidel CC. 1932. Studies of living nerves. I. The movements of individual sheath cells and nerve sprouts correlated with the process of myelin-sheath formation in amphibian larvae. J Exp Zool 61:279–331. doi:10.1002/jez.1400610206

Sukumaran SK, Lewandowski BC, Qin Y, Kotha R, Bachmanov AA, Margolskee RF. 2017. Whole transcriptome profiling of taste bud cells. Sci Rep 7:7595. doi:10.1038/s41598-017-07746-z

Takahashi S, Ishida A, Kubo A, Kawasaki H, Ochiai S, Nakayama M, Koseki H, Amagai M, Okada T. 2019. Homeostatic pruning and activity of epidermal nerves are dysregulated in barrier-impaired skin during chronic itch development. Sci Rep 9:8625. doi:10.1038/s41598-019-44866-0

Tang JY, Mackay-Wiggan JM, Aszterbaum M, Yauch RL, Lindgren J, Chang K, Coppola C, Chanana AM, Marji J, Bickers DR, Epstein EH. 2012. Inhibiting the Hedgehog Pathway in Patients with the Basal-Cell Nevus Syndrome. N Engl J Med 366:2180–2188. doi:10.1056/NEJMoa1113538

Trachtenberg JT, Chen BE, Knott GW, Feng G, Sanes JR, Welker E, Svoboda K. 2002. Long-term in vivo imaging of experience-dependent synaptic plasticity in adult cortex. Nature 420:788–794. doi:10.1038/nature01273

Treffy RW, Collins D, Hoshino N, Ton S, Katsevman GA, Oleksiak M, Runge EM, Cho D, Russo M, Spec A, Gomulka J, Henkemeyer M, Rochlin MW. 2016. Ephrin-B/EphB Signaling Is Required for Normal Innervation of Lingual Gustatory Papillae. Dev Neurosci 38:124–138. doi:10.1159/000444748

Vandenbeuch A, Larson ED, Anderson CB, Smith SA, Ford AP, Finger TE, Kinnamon SC. 2015. Postsynaptic P2X3-containing receptors in gustatory nerve fibres mediate responses to all taste qualities in mice: Role of P2X3-containing receptors in taste. J Physiol 593:1113–1125. doi:10.1113/jphysiol.2014.281014

Verzé L, Paraninfo A, Ramieri G, Viglietti-Panzica C, Panzica GC. 1999. Immunocytochemical evidence of plasticity in the nervous structures of the rat lower lip. Cell Tissue Res 297:203–211. doi:10.1007/s004410051348

Walsh MK, Lichtman JW. 2003. In Vivo Time-Lapse Imaging of Synaptic Takeover Associated with Naturally Occurring Synapse Elimination. Neuron 37:67–73. doi:10.1016/S0896-6273(02)01142-X

Wilson CE, Lasher RS, Yang R, Dzowo Y, Kinnamon JC, Finger TE. 2022. Taste Bud Connectome: Implications for Taste Information Processing. J Neurosci 42:804. doi:10.1523/JNEUROSCI.0838-21.2021

Wu A, Dvoryanchikov G, Pereira E, Chaudhari N, Roper SD. 2015. Breadth of tuning in taste afferent neurons varies with stimulus strength. Nat Commun 6:1–11.

Yarmolinsky DA, Zuker CS, Ryba NJP. 2009. Common Sense about Taste: From Mammals to Insects. Cell 139:234–244. doi:10.1016/j.cell.2009.10.001

Yee CL, Jones KR, Finger TE. 2003. Brain-derived neurotrophic factor is present in adult mouse taste cells with synapses. J Comp Neurol 459:15–24. doi:10.1002/cne.10589

Zaidi FN, Cicchini V, Kaufman D, Ko E, Ko A, Van Tassel H, Whitehead MC. 2016. Innervation of taste buds revealed with Brainbow-labeling in mouse. J Anat 229:778–790. doi:10.1111/joa.12527

Zaidi FN, Whitehead MC. 2006. Discrete Innervation of Murine Taste Buds by Peripheral Taste Neurons. J Neurosci 26:8243–8253. doi:10.1523/JNEUROSCI.5142-05.2006

Zeng H, Sanes JR. 2017. Neuronal cell-type classification: challenges, opportunities and the path forward. Nat Rev Neurosci 18:530–546. doi:10.1038/nrn.2017.85

Zhang J, Jin H, Zhang W, Ding C, O’Keeffe S, Ye M, Zuker CS. 2019. Sour Sensing from the Tongue to the Brain. Cell 179:392-402.e15. doi:10.1016/j.cell.2019.08.031

